# Machine Learning Identifies Complicated Sepsis Course and Subsequent Mortality Based on 20 Genes in Peripheral Blood Immune Cells at 24 Hours post ICU admission

**DOI:** 10.1101/2020.06.14.150664

**Authors:** Shayantan Banerjee, Akram Mohammed, Hector R. Wong, Nades Palaniyar, Rishikesan Kamaleswaran

## Abstract

A complicated clinical course for critically ill patients admitted to the ICU usually includes multiorgan dysfunction and subsequent death. Owning to the heterogeneity, complexity, and unpredictability of the disease progression, patient care is challenging. Identifying the predictors of complicated courses and subsequent mortality at the early stages of the disease and recognizing the trajectory of the disease from the vast array of longitudinal quantitative clinical data is difficult. Therefore, we attempted to identify novel early biomarkers and train the artificial intelligence systems to recognize the disease trajectories and subsequent clinical outcomes. Using the gene expression profile of peripheral blood cells obtained within 24 hours of PICU admission and numerous clinical data from 228 septic patients from pediatric ICU, we identified 20 differentially expressed genes that were predictive of complicated course outcomes and developed a new machine learning model. After 5-fold cross-validation with ten iterations, the overall mean area under the curve reached 0.82. Using the same set of genes, we further achieved an overall area under the curve of 0.72 when tested on an external validation set. This model was highly effective in identifying the clinical trajectories of the patients and mortality. Artificial intelligence systems identified eight out of twenty novel genetic markers *SDC4, CLEC5A, TCN1, MS4A3, HCAR3, OLAH, PLCB1* and *NLRP1* that help to predict sepsis severity or mortality. The discovery of eight novel genetic biomarkers related to the overactive innate immune system and neutrophils functions, and a new predictive machine learning method provides options to effectively recognize sepsis trajectories, modify real-time treatment options, improve prognosis, and patient survival.

**Research in Context:** *Evidence before this study:* Transcriptomic biomarkers have long been explored as potential means of earlier disease endotyping. Much of the existing literature has however focused on mortality and discrete outcomes. Additionally, much of prior work in this area has been developed on statistical methods, while recent means of selecting features have not been sufficiently explored.

*Added value of this study:* In this study, we developed a robust machine learning based model for identifying novel biomarkers of complicated disease courses. We found 20 highly stable genes that predict disease complexity with an average derivation AUROC of 0.82 and validation AUROC of 0.72 within critically ill children, using peripheral blood collected within 24 hrs of ICU admission.

*Implications of all the available evidence:* Earlier identification of disease complexity can inform care management and targeted therapy. Therefore, the 20 gene candidates identified by our rigorous approach, can be used to identify, early in their ICU stay, patients who may ultimately develop significant organ dysfunction and complex care management.

## Introduction

Critically ill patients are admitted to the Intensive Care Unit (ICU) for complex and dynamic care, preserving organ function and improving outcomes in otherwise dire situations. Among patients with complicated disease courses, septic patients represent a significant component (1). Sepsis is a life-threatening organ dysfunction caused by the overactive immune response to bacterial infection, often of pulmonary origin (2). Sepsis may have contributed to 20% of all global deaths in 2019 (3). Sepsis consists of a heterogeneous mix of phenotypes (4, 5), various degrees of disease complexities, and trajectories leading to recovery or death (6, 7). Different strategies have been pursued predicting deterioration (8–10) and managing patients with sepsis in critical care units (11) using physiological, clinical, and biomarker parameters. However, due to the heterogeneous nature of patients presenting to the ICU, and the diverse disease course that follows, it has been difficult to identify generalized models of disease (12).

Statistical and machine learning methods have been developed to successfully utilize the multi-omics data for biomarker discovery for predicting survival from sepsis (13). Wong et al. (14) identified 12 biomarkers collectively called the Pediatric Sepsis Biomarker Risk Model (PERSEVERE) class genes. Further analyses resulted in the identification of 18 additional genes consisting of the PERSEVERE XP set (15). Mohammed et al. (16) identified 53 differentially expressed genes involved mostly in immune response and chemokine activity from expression data collected from patients admitted to the PICU within 24 hours of their admission. Sweeney et al. (17) analyzed the results obtained from three independent scientific groups that developed mortality prediction models and identified additional subgroups of genes. While much has been studied about the risk for mortality, there is a dearth of machine learning approaches to predict disease trajectory, including complicated disease courses and poor clinical outcomes (18). Early identification of disease trajectory, including complicated disease courses, defined as persistence of 2 or more organ failures by day 7 or death by day 28, can aid in clinical management and targeted therapies to manage severe outcomes. Hence, there is a need to identify these biomarkers and build novel machine learning models to identify complex disease courses from plasma samples collected close to the time of ICU admission.

In this work, we used peripheral blood cell gene expression data (sampled within 24 hours of the onset of sepsis; 228 pediatric ICU patients) and analyzed them using multiple statistical and machine learning methods to identify novel markers of complicated disease course. We validated these features by calculating their overlap with the well-established sepsis mortality predicting genes, conducting the functional gene-set enrichment and pathway analyses and testing them on an external validation dataset.

## Materials and Methods

### Data collection

The pediatric sepsis dataset GSE66099 (19) contains the gene expression profiles extracted from the peripheral blood samples of patients who were admitted to the PICU during the first 24 hours of admission. The microarray dataset was obtained from the Affymetrix GPL570 platform, which was submitted by Wong et al. on February 19, 2015, and last updated on March 25, 2019. We considered a complicated course as the primary outcome variable. It was defined as death by 28 days or persistence >=2 organ failures at day 7 of septic shock (20). This dataset was used to train our model and derive the top gene features.

Due to the lack of available external data that encodes for complicated courses, we used the closest surrogate microarray gene expression dataset GSE54514 (21), which provides whole blood samples collected up to 5 days after ICU admission. In this cohort, we defined the complicated presentation as patients with an APACHE II score >=25 observed within 24 hours of admission to the ICU.

### Normalization and background correction

Technical variations in the gene expression data were eliminated using the Affy package (22) from R. This also helped remove the background noise. The Robust Multi-Average parameter method using ‘gcrma’ package (33) from R was used to normalize data and perform background correction. The resulting variation was removed using the comBat (24) and limma linear fit model.

### Probe to gene mapping, identification of differentially expressed genes

For the two microarray datasets GSE66099 and GSE54514, the Affymetrix probes were matched to gene symbols using the Affymetrix database (hgu133plus2.db) and Illumina HumanHT12v4 annotation database (illuminaHumanv4.db), respectively. In order to detect the expression values of genes with multiple probes, we averaged the expression for multiple probes that matched to the same gene. The limma package (25) from R was used to perform the differential gene expression analysis with a Benjamini-Hochberg (BH) correction.

### Functional enrichment analysis

We used the clusterProfiler package (26) in R to find enriched biological processes (BP), cellular components (CC), and molecular function (MF) terms. A plot displaying the enriched terms was drawn using the enrichplot package (27) in R. The STRING database v11 (28) was used to find significantly enriched Reactome and KEGG pathways.

### Statistical analysis

Affymetrix data download and gene mapping were done using the R Affy and Bioconductor packages. A Fisher exact test was performed to determine the statistical significance among the functional enrichment terms. Benjamini Hochberg multiple test correction method was used to calculate the DEGs. Heatmap for the DEGs was generated using the complexHeatmap package in R. Volcano plot and the MA plot were generated using the plotMA and volcanoplot of the limma package.

### Feature selection methods

The processed microarray derivation dataset had 20,174 genes for 228 samples. Our next objective was to reduce the dimensionality of the dataset by removing redundant features. To do that we used three commonly used feature selection techniques including random forest-based feature importance, LASSO, and Minimum Redundancy and Maximum Relevance (MRMR). The features generated by each of the above three methods were pooled together into one aggregated feature set and the list of 17 differentially expressed genes were also added to that list. The Pediatric Risk of Mortality Score (PRISM) for each patient was computed and included in the model (29). For external validation, we used LASSO based feature selection technique to select the top features for further analysis.

### Classification models

Imbalanced data can negatively affect learning algorithms, hence, in this study, we experimented with both oversampling and undersampling techniques to balance the training data. We then developed machine learning algorithms using two tree-based classifiers and logistic regression. The two tree-based classifiers were the balanced random forest and extra trees classifier. The balanced random forest classifier balances data by randomly undersampling each bootstrapped sample. The extra trees classifier is a meta estimator that fits a user-defined number of randomized trees (base models) on different sub-samples of the data and then combines the predictive power of these models into one optimal model.

### Model selection and tuning

The learning process that was adopted in the manuscript is illustrated in Figure 1(A-C). First, the dataset was randomly partitioned in a stratified fashion into five equal subsets (Figure 1A). Four of the five subsets were combined into one training set. In each training phase, we performed (1) feature selection (Figure 1B): The pooled list of features using the three feature reduction approaches (described in the previous section), along with statistical filtering by DEG were obtained. Finally, Recursive Feature Elimination (RFE) was performed on the pooled features to remove redundancies and arrive at a subset of the most important genes. 2) Hyperparameter tuning was performed using a cross-validated grid search technique over a parameter grid using the AUROC metric as the scoring function. 3) A classifier was trained using the hyperparameters, and the corresponding prediction scores were obtained on the hold-out test set (Figure 1C). We repeated the entire process ten times, resulting in 50 unique train and test splits. The reported performances obtained during each run were averaged and the mean scores were reported. We assessed the performance of our classifiers using four widely used performance metrics: Sensitivity, Specificity, Mathew’s correlation coefficient, and Area under the ROC curve (AUROC). Accuracy and F1 score are widely used performance metrics that can show overoptimistic inflated results for imbalanced data and hence were not included in our study. To study the predictive power of our top 20 gene features from the stability analysis, we performed validation on an independent test set. In the first step, we applied different undersampling and oversampling techniques on the derivation dataset. Then we performed LASSO based feature selection to remove redundant features. The hyperparameters of the classifier were tuned using the 5-fold cross-validation grid search technique over a parameter grid using the AUROC metric as the scoring function. The tuned classifier was then trained on the derivation set and the corresponding prediction scores were obtained on the hold-out validation set (GSE54514). To further fine-tune the classifiers, instead of using the default classification threshold of 0.5, we experimented with different thresholds between 0 and 1 with step sizes of 0.001. This is particularly important for an imbalanced classification problem like ours and must be taken into account to make sure that the minority class examples are predicted correctly. The same set of performance metrics were used to assess the quality of the validation classifier as stated in the previous section.

**Figure 1:**
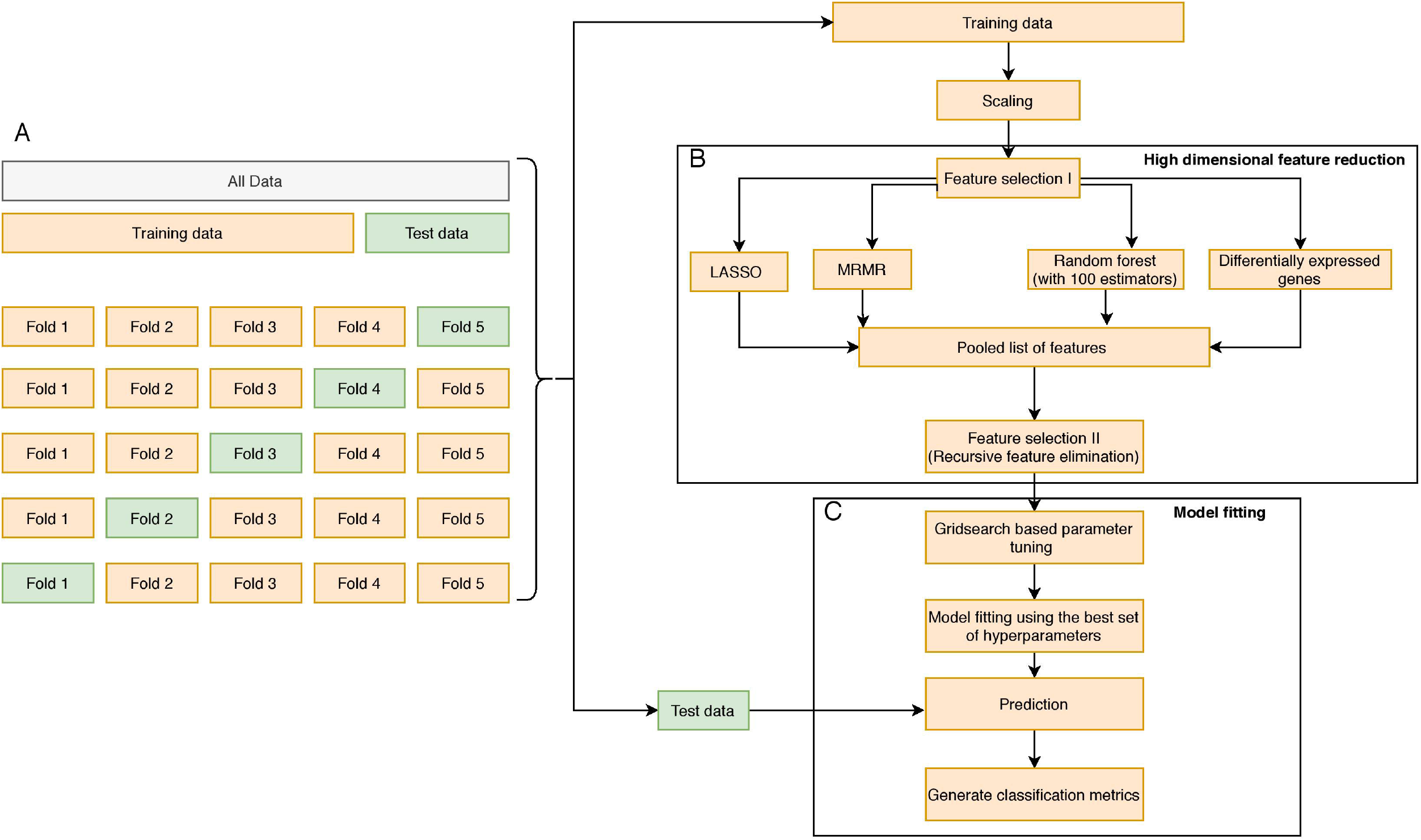
The overall methodology design is illustrated in this figure. (A) The initial data is aggregated, normalized, corrected for batch normalization, and separated into even chunks using k-fold cross-validation (CV). In our pipeline, we used k=5. (B) The training chunks of the CV are used for model development; the data analysis pipeline follows the Complete Cross-Validation (CCC) approach defined by Simon et al. (51). In addition to DEG, we apply three other feature selection methods to generate a pool of candidate genes. We then apply a wrapper method, namely the RFE to arrive on the most predictive genes. (C) The genes selected by the RFE method are then used to develop a predictive model. The model is then evaluated on the test fold of the CV. This process is repeated for the remaining training and test folds. Finally, the entire 5-fold CV is repeated ten times to generate a total of 50 iterations, and the top predictors from (B) are saved and analyzed to generate a normalization score, which is a measure of how often a gene appears as a top predictor across each of the 50 iterations.

### Distribution of features

To study the class-wise distributional differences between the features, we conducted the two-sample Kolmogorov-Smirnov test using scipy stats module. The distplot function from the seaborn package was used to visualize the class-wise Gaussian kernel density estimate of each of the features.

## Results - Derivation dataset

### Patient characteristics

Out of the 229 pediatric septic patients, 52 had a complicated course outcome. One patient was excluded due to missing the outcome variable. The age of the cohort was 3.81±3.42 (mean±SD) years. Out of the 228 patients in the cohort, 18 patients met the criteria for sepsis, 30 for SIRS, and 180 for septic shock. Males constitute the majority of the dataset (139; 61%; *P*=.00093). In the 52 complicated course cohort, 31 (59.6%; *P*=.16) were male; whereas in the non-complicated course group, 108 (61.36 %; *P*=.0026) were male. Of the 52 with complicated courses, 28 (53.8%; *P*=.58) died. The clinical characteristics of all patients with a complicated or uncomplicated course outcome are provided in Supplementary Table 1a. Microbiologic results are shown in Supplementary Table 3. The most frequently identified organisms were *Staph aureus* (in 22 patients; 9.65%), *Pneumococcus* (in 18 patients; 7.9%), *Group A Strep* (in 14 patients; 6.14%), and *Klebsiella* (in 11 patients; 4.82%).

### Gene clustering

Figure 2 (A-C) shows the average gene expression of samples before and after normalization and represents the batch effects after adjusting for longitudinal processing of the microarrays. Figure 3(A) illustrates the heatmap of 17 DEGs (absolute fold change > 1 and adjusted p-value <0.1) across the complicated vs uncomplicated groups. The samples are represented on the vertical axis and the genes are on the horizontal axis. The log2 gene expression values of the DEGs were used to construct the heatmap. Red and green represent the upregulated and downregulated genes, respectively. Both the samples and genes were clustered to give a better idea of the groups formed using the expression values. Two horizontal bars show annotations for the complicated course outcomes and mortality.

**Figure 2:**
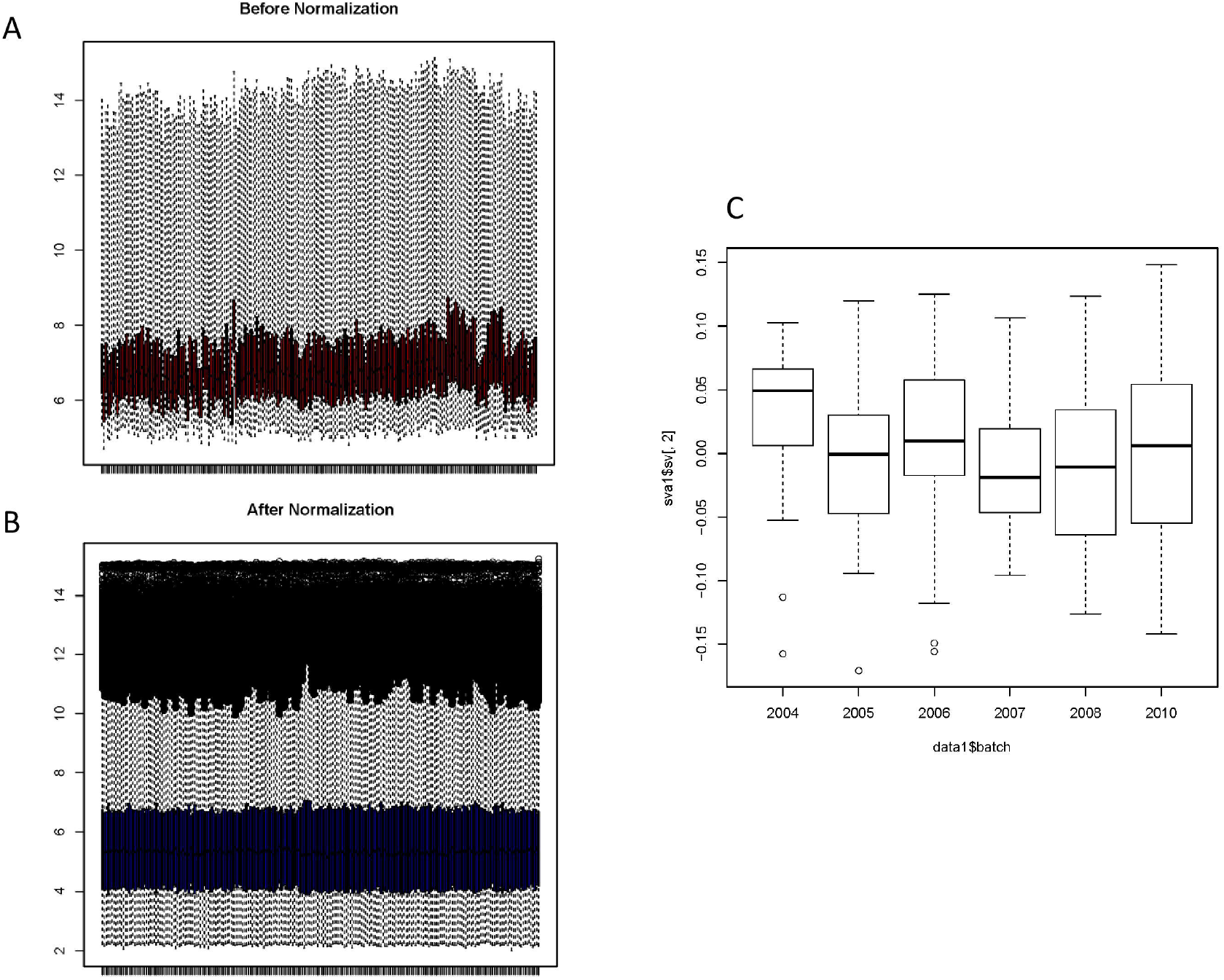
Preprocessing of microarray data: (A) and (B) Average gene expression values before normalization and after normalization. The x-axis represents the samples, and the y-axis represents the gene expression values. According to the figures, the average expression values of the samples were more stable and consistent after normalization and suitable for analysis. (C) One of the most well-known sources of variation in gene expression studies is batch effects when samples are processed during different time points or by different groups of people. We removed the batch effect from the data due to the microarray experiments being conducted over multiple years. In the given figure, the second SVA component ordered by date before batch effect correction displays non-uniformity in the expression values.

**Figure 3:**
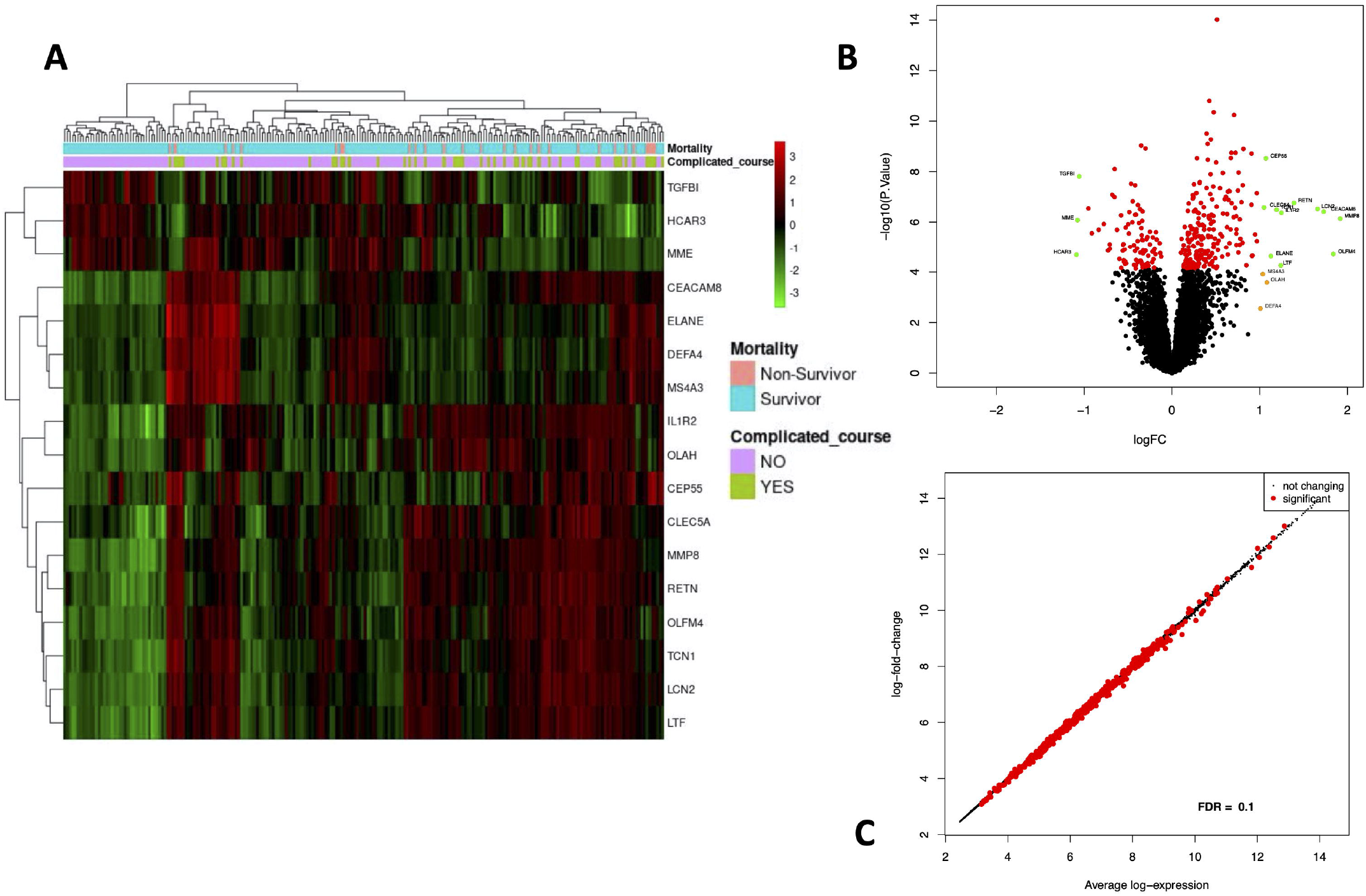
Differential gene expression analysis results are compiled in this figure. (A) Heatmap representing differentially expressed genes between complicated and uncomplicated course groups with annotations. The color and intensity of the boxes represent changes in gene expression. Red represents upregulated genes and green represents downregulated genes. The horizontal bars show annotations for complicated course outcomes and mortality and are useful for interpreting the sample-wise clusters formed using the expression measurements. (B) Volcano plot of differentially expressed genes in complicated and uncomplicated course outcomes. A volcano plot helps us to assess the adjusted p values (significance), and the log fold changes (biological difference) of differential expression for the given list of genes at the same time. (C) An MA plot is a 2D scatter plot (each dot representing a gene) that represents log fold change vs mean expression across two different conditions. All the significantly differentially expressed genes (FDR cutoff =0.1) are colored in red and the genes without significant gene expression differences are colored in black.

Three prominent sample clusters were observed from the heatmap. The first cluster contained none of the 52 complicated course patients, while the second cluster contained nine (17%) of the 52 complicated course patients and the third contained 43 (83%) of the complicated course patients. A similar cluster trend is noticed among survivors/non-survivors. None of the non-survivors were in the first cluster, four (14%) out of 28 non-survivors were in the second cluster and the remaining 24 (86%) were in the third cluster.

Genes were also clustered into two distinct groups of upregulated and downregulated genes. *MME, TGFBI*, and *HCAR3* together formed one cluster of downregulated genes while the rest were in the upregulated category. *LTF* and *IL1R2* were the most highly expressed genes, whereas *CEP55*, and *MS4A3* were the least expressed genes. *MMP8* was the most upregulated gene while *HCAR3* was the most downregulated gene.

Based on an adjusted p-value cutoff < 0.1, a total of 1269 differentially expressed genes (DEGs) from 20,174 were found between the complicated and uncomplicated group of patients, including 808 upregulated genes and 461 downregulated genes. The DEGs with a fold change of at least one (n=17) are shown in Supplementary Table 2. (For the complete list, refer to Supplementary Table 4). Figure 3B shows the volcano plot of significantly different genes. Genes represented by green dots are logFoldChange>1 and adjusted p-value <0.1; red dot represents genes with adjusted p-value < 0.1 but logFoldChange<1; whereas black represent genes with logFoldChange<1 and adjusted p-value >0.1. Also, in Figure 3C, an MA-plot displays the log fold change between complicated course and non-complicated course samples as a function of the average expression level across all samples. Red dots are relatively larger than the black ones. The dataset included a total of 30 uncomplicated disease course patients who met the SIRS criteria when excluding these patients from our analysis we observed five genes that achieved robust normalization scores but did not meet the DEG cutoffs. Those genes were: *TGFBI, DEFA4, CEP55, MME, and OLAH*.

### Functional enrichment analysis of the DEGs

The top 17 most significantly differentially expressed genes (absolute log fold change > 1 and adjusted p-value <0.1) were analyzed. A plot displaying the functional analysis is shown in Figure 4. Hematopoietic cell lineage (hsa04640) was the most enriched KEGG pathway; neutrophil degranulation (HSA-6798695), immune system (HSA-168256) and antimicrobial peptides (HSA-6803157) were some of the most enriched Reactome pathways. Gene Ontology (GO) analysis of the DEGs revealed that neutrophil degranulation (GO:0043312), neutrophil activation involved in immune response (GO:0002283), neutrophil activation (GO:0042119) and neutrophil-mediated immunity (GO: 0002446) as some of the top enriched biological processes. Serine-type endopeptidase activity (GO:0004252), endopeptidase activity (GO: 0004175), and serine-type peptidase activity (GO:0008236) were some of the most enriched molecular functions; Specific granule (GO:0042581) and specific granule lumen (GO:0035580) were some of the most enriched cellular components.

**Figure 4:**
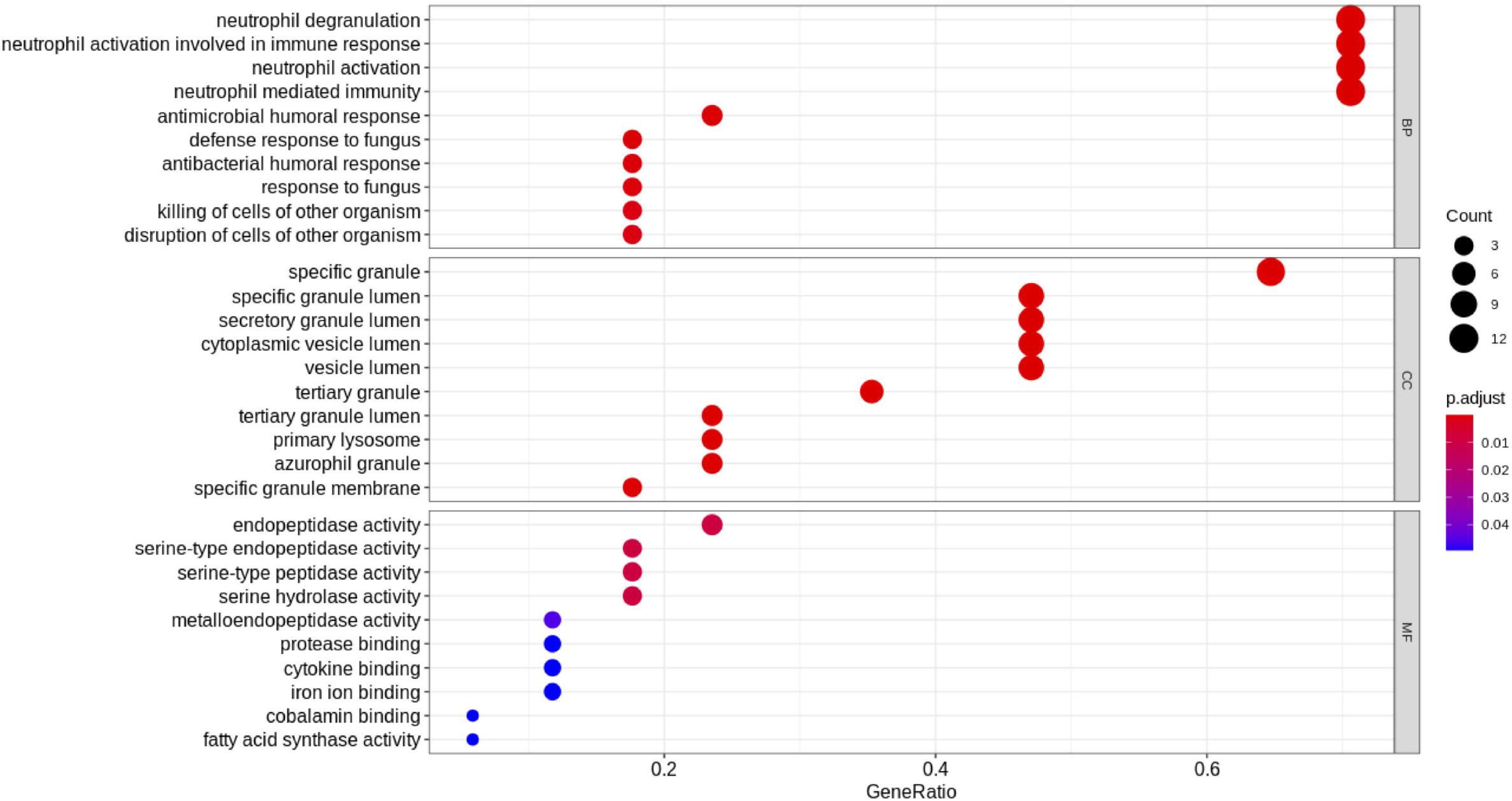
Functional analysis of the 17 DEGs. The size of the dot represents the number of genes associated with that enriched term and the color of the dot represents the significance of the terms (more significant terms being redder).

We also performed functional analysis on the top predictors from our machine learning analysis which included three additional genes *(PLCB1, NLRP1 and SDC4)* apart from the DEGs (Table 3). The top enriched biological process term was neutrophil degranulation (GO:0043312) similar to the results from our previous functional analysis with the DEGs. Some of the newer enriched terms included cell activation (GO:0001775), immune response (GO: 0006955), immune system process (GO:0002376), and response to stimulus (GO: 0050896). It is well known that immunosuppression is a hallmark of sepsis and the top predictors capable of distinguishing between complicated and uncomplicated course patients show enrichment in the immune response process. Neutrophil degranulation (HSA-6798695), innate immune system (HSA-168249) were among the top enriched REACTOME pathways. The innate immune system is activated as the first response to an infection before the adaptive immune system. Most of the clinical phenotypes of sepsis can be attributed to the innate immune response. Interestingly, the adaptive immune response was not one of the enriched pathways for our list of genes which might warrant further investigation.

### Predictive performance

The best performance in terms of overall mean AUROC (=0.823) was obtained using an oversampling technique, SMOTE, and BRF classifier pair. This model also gave the best mean specificity (=0.885) and the best mean MCC (=0.445). The best mean sensitivity (=0.792) was obtained using an undersampling technique, REDN, and the BRF classifier. The ROC curve of the best performing model is shown in figure 5 and the results for the best performing classifiers is shown in Table 1. The orange-colored curve is the mean ROC curve obtained after averaging the results from different runs of the repeated 5-fold CV. The list of the top features that were consistently chosen across different folds of the cross-validation of the top-performing model and their overlap with previously published predictor genes in sepsis is shown in Table 3. The ‘normalized score’ represents the fraction of times the given predictor was consistently chosen over all experiments, representing the stability (30) of these genes as possible predictors of a complicated course.

**Figure 5:**
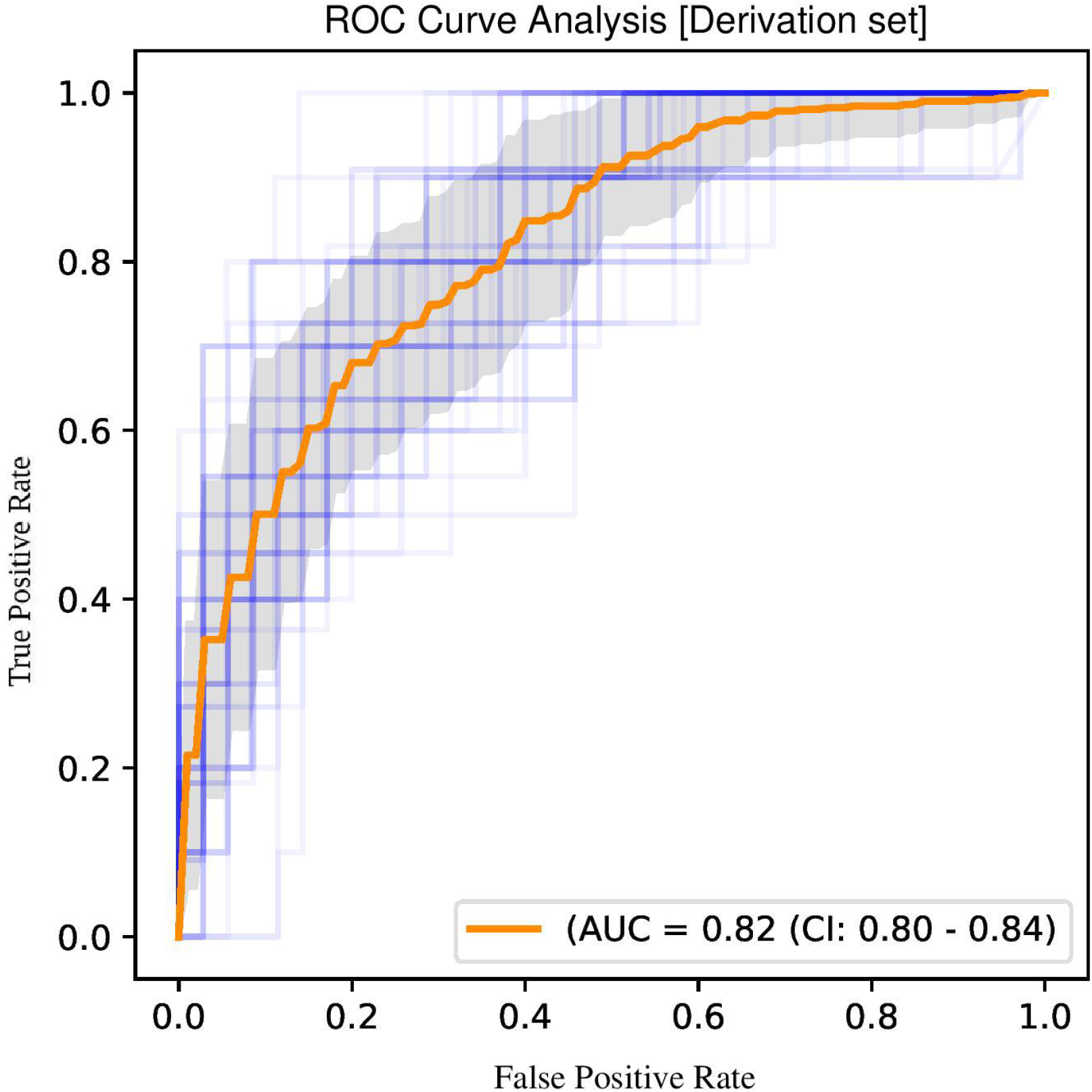
A ROC plot illustrates the performance of a binary classifier at different classification thresholds usually featuring a false positive rate (1-Specificity) on the x-axis and true positive rate (Sensitivity) on the y-axis. The top left corner is an ideal point with a false positive rate of zero and a true positive rate of one. The area under the curve (or AUC) denotes the probability that a randomly chosen positive instance is ranked higher than a randomly chosen negative one by our classifier. An AUC of zero means that the classifier is predicting the positive class as negative and vice versa, while an AUC of one denotes perfect separability. In the above figure, we denote the ROC plots generated from the different cross-validation experiments along with the mean area under the curve (in orange). The variance of the curve (shaded part) roughly shows how the output from our best performing model is affected by changes in the training data.

**Table 1:**
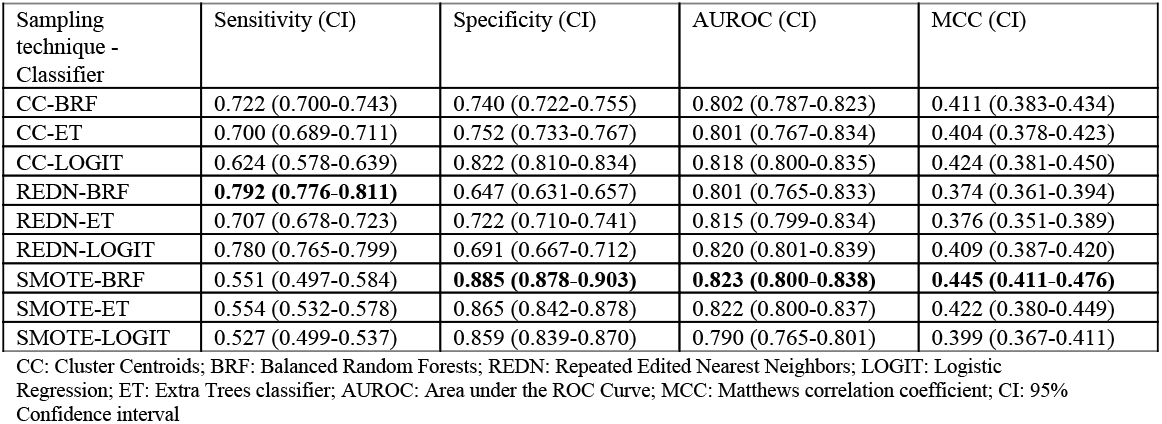
Performance measure for various sampling technique and classifier pairs (Derivation set)

| Sampling technique - Classifier | Sensitivity (CI) | Specificity (CI) | AUROC (CI) | MCC (CI) |
| --- | --- | --- | --- | --- |
| CC-BRF | 0.722 (0.700-0.743) | 0.740 (0.722-0.755) | 0.802 (0.787-0.823) | 0.411 (0.383-0.434) |
| CC-ET | 0.700 (0.689-0.711) | 0.752 (0.733-0.767) | 0.801 (0.767-0.834) | 0.404 (0.378-0.423) |
| CC-LOGIT | 0.624 (0.578-0.639) | 0.822 (0.810-0.834) | 0.818 (0.800-0.835) | 0.424 (0.381-0.450) |
| REDN-BRF | **0.792 (0.776-0.811)** | 0.647 (0.631-0.657) | 0.801 (0.765-0.833) | 0.374 (0.361-0.394) |
| REDN-ET | 0.707 (0.678-0.723) | 0.722 (0.710-0.741) | 0.815 (0.799-0.834) | 0.376 (0.351-0.389) |
| REDN-LOGIT | 0.780 (0.765-0.799) | 0.691 (0.667-0.712) | 0.820 (0.801-0.839) | 0.409 (0.387-0.420) |
| SMOTE-BRF | 0.551 (0.497-0.584) | **0.885 (0.878-0.903)** | **0.823 (0.800-0.838)** | **0.445 (0.411-0.476)** |
| SMOTE-ET | 0.554 (0.532-0.578) | 0.865 (0.842-0.878) | 0.822 (0.800-0.837) | 0.422 (0.380-0.449) |
| SMOTE-LOGIT | 0.527 (0.499-0.537) | 0.859 (0.839-0.870) | 0.790 (0.765-0.801) | 0.399 (0.367-0.411) |

CC: Cluster Centroids; BRF: Balanced Random Forests; REDN: Repeated Edited Nearest Neighbors; LOGIT: Logistic Regression; ET: Extra Trees classifier; AUROC: Area under the ROC Curve; MCC: Matthews correlation coefficient; CI: 95% Confidence interval

**Table 2:** Performance measure for various sampling technique and classifier pairs (Validation set)

| Sampling technique - Classifier | Sensitivity (95% CI) | Specificity (95% CI) | AUROC (95% CI) | MCC (95% CI) |
| --- | --- | --- | --- | --- |
| IHT-BRF | 0.667 (0.662-0.672) | 0.740 (0.732-0.746) | 0.703 (0.691-0.708) | 0.349 (0.335-0.355) |
| IHT-ET | **0.704 (0.688-0.709)** | 0.700 (0.687-0.712) | 0.701 (0.696-0.709) | 0.339 (0.330-0.345) |
| IHT- BRF+ET | 0.629 (0.619-0.635) | **0.800 (0.789-0.810)** | 0.714 (0.709-0.718) | 0.386 (0.379-0.390) |
| IHT- ET+GB | 0.629 (0.618-0.634) | 0.790 (0.780-0.808) | 0.709 (0.701-0.712) | 0.374 (0.370-0.380) |
| IHT- BRF+GB+ET | 0.629 (0.619-0.635) | **0.800 (0.788-0.809)** | 0.715 (0.710-0.720) | 0.387 (0.378-0.390) |
| REDN-BRF | 0.670 (0.652-0.681) | 0.780 (0.758-0.811) | **0.723 (0.719-0.727)** | **0.395 (0.380-0.400)** |
| REDN-EE | 0.629 (0.619-0.635) | 0.770 (0.741-0.779) | 0.700 (0.682-0.709) | 0.352 (0.340-0.360) |
| REDN-BRF+LOGIT | 0.670 (0.652-0.682) | 0.770 (0.741-0.779) | 0.718 (0.710-0.721) | 0.382 (0.375-0.391) |
| REDN- BRF+XGB | 0.670 (0.652-0.682) | 0.740 (0.731-0.749) | 0.703 (0.691-0.709) | 0.350 (0.341-0.359) |

IHT: Instance Hardness Threshold; BRF: Balanced Random Forests; REDN: Repeated Edited Nearest Neighbors; LOGIT: Logistic Regression; ET: Extra Trees classifier; EE: Easy Ensemble; GB: Gradient Boosting; XGB: XGBoost; AUROC: Area under the ROC Curve; MCC: Matthews correlation coefficient

**Table 3:**
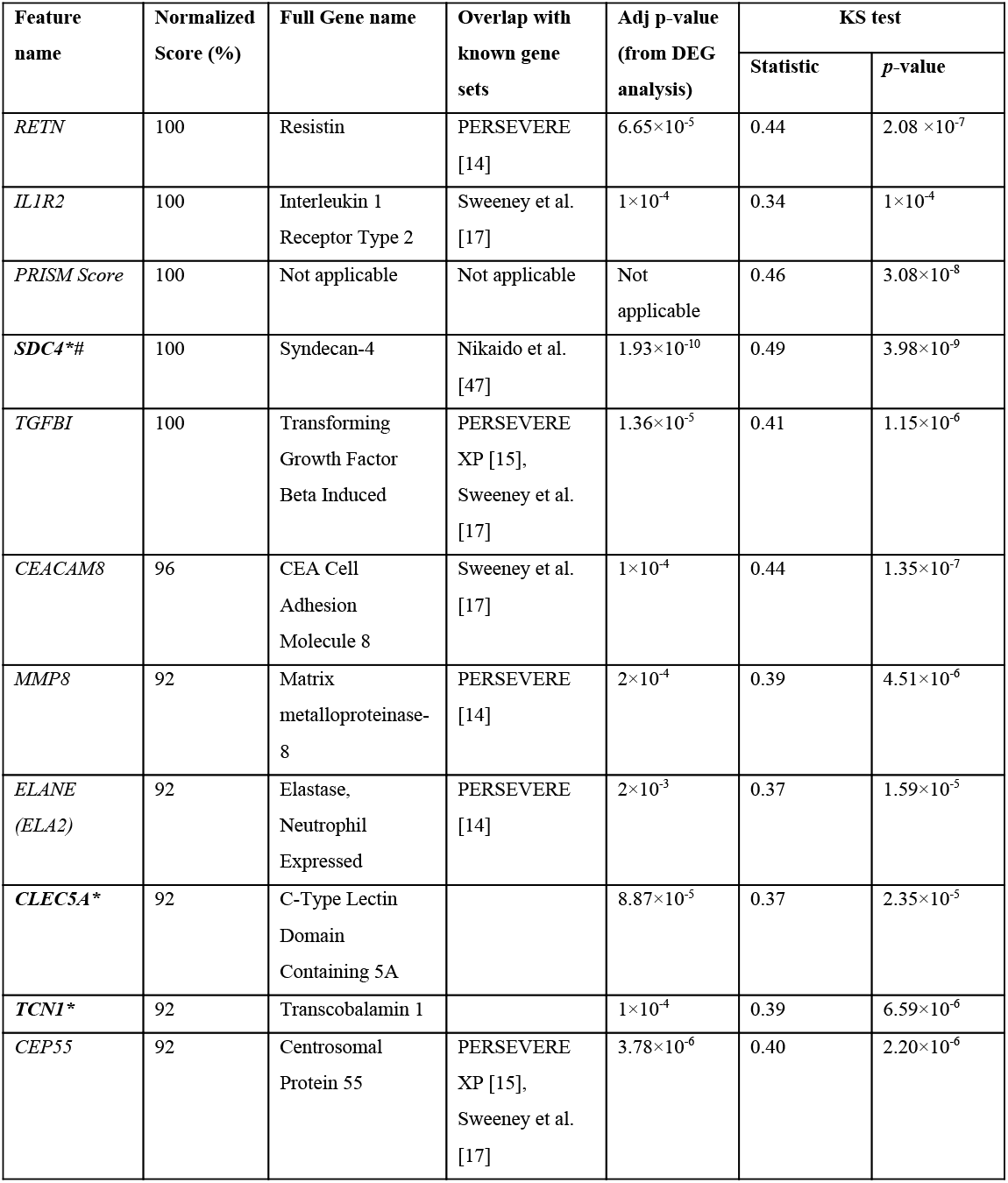
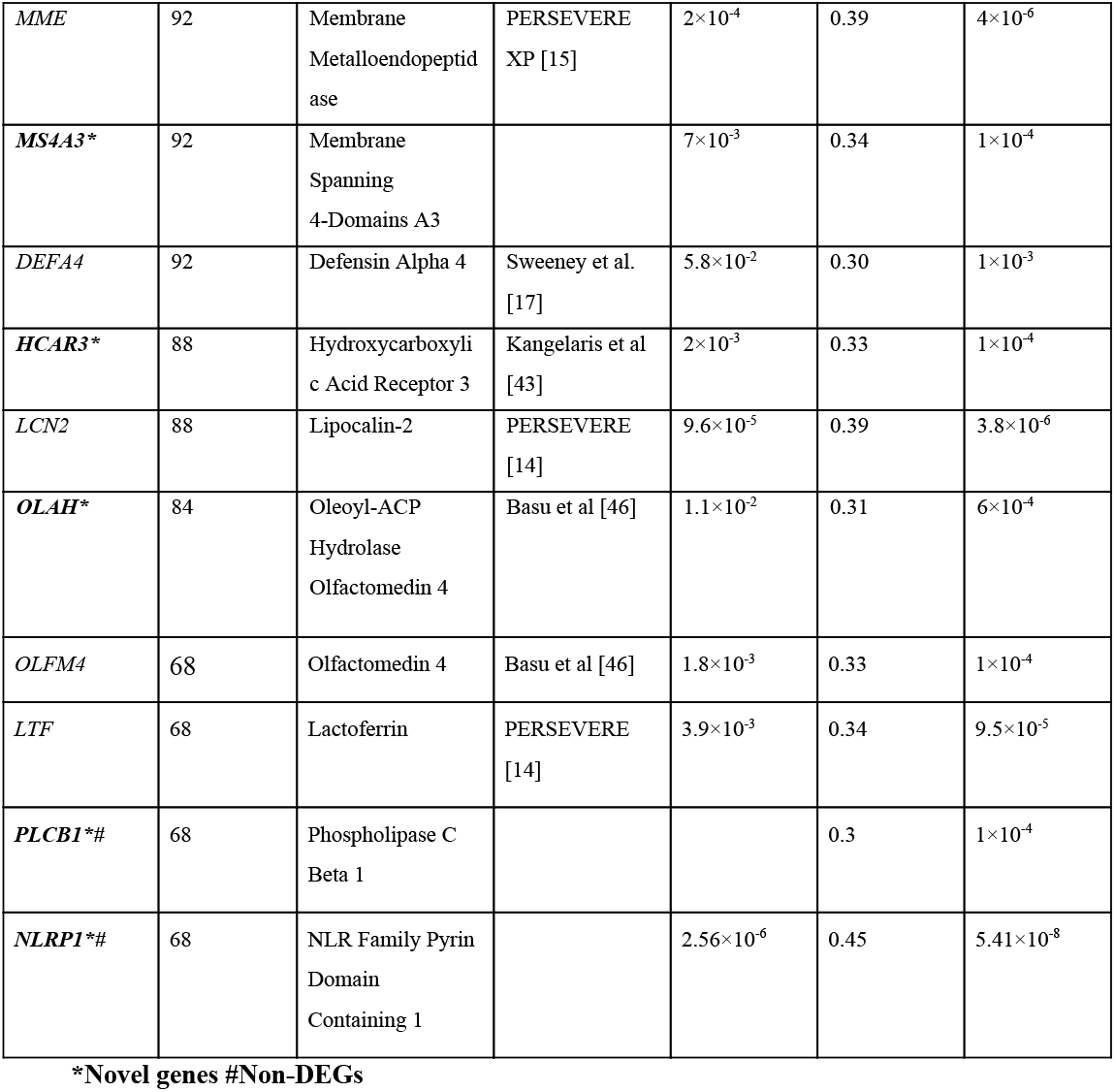
Top consistently chosen features across folds

## Results - Validation dataset

### Patient characteristics

Out of the 127 septic patients, 27 had a complicated course outcome. The age of the cohort was 59.13±15.99 (mean±SD). Females constitute the majority of the dataset (75; 59%; *P*=.11). In the 27 complicated course patients, 14 (51.85%;*P*=0.02) were females; whereas, in the non-complicated course group, 61 (61%; *P*=0.14) were females. Of the 27 with the complicated course, 17 (63%; *P*=0.16) died. The clinical characteristics of all patients with a complicated or uncomplicated course outcome are provided in Supplementary Table 1b.

### Predictive performance

The best performance in terms of overall AUROC (=0.7233) was obtained using an undersampling technique, REDN, and BRF classifier pair. This model also gave the best MCC (0.395). The best specificity (0.80) was obtained using another undersampling technique, IHT and a stacked classifier containing BRF, GB and ET. The best sensitivity (0.70) was obtained using the sampling technique, IHT and ET classifier pair. The ROC curve for the top-performing models is shown in figure 6 and the classification metrics for the same are shown in Table 2. For the undersampling technique REDN, the list of features that were selected using the LASSO based feature reduction technique were *MMP8, CEACAM8, LCN2, RETN, CLEC5A, TGFBI, CEP55, MME, OLAH, SDC4* and for IHT the list of features that were selected was *OLFM4, CEACAM8, LCN2, RETN, ELANE, HCAR3, IL1R2, CLEC5A, MS4A3, TGFBI, DEFA4, CEP55, MME, SDC4, PLCB1, NLRP1*. The list of tuned hyperparameters for the different sampling-classifier pairs that gave the best results is attached in Supplementary Table 5a and the details of the other classifiers that were experimented with are attached in Supplementary table 5b.

**Figure 6:**
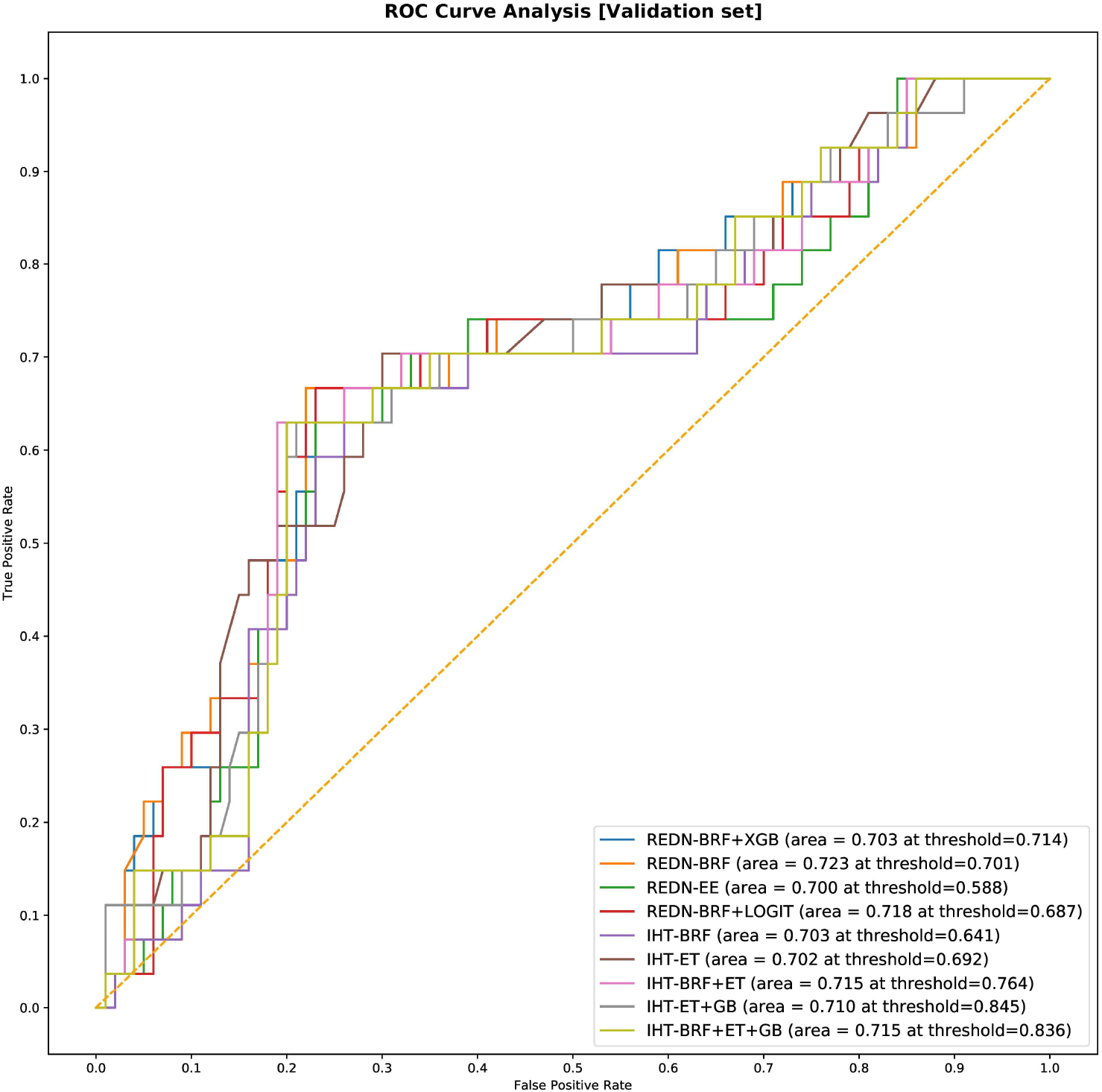
A combined ROC plot illustrating the validation set performance of the different binary classifiers is shown in this figure. Due to the inherent imbalance in the training data (GSE66099) we tried different classification thresholds for the tuned classifier and reported the ones that gave the best classification performances. The list of top performing classifiers included both individual and stacked classifiers. The legend displays the names of the classifiers, the AUROC and the classification thresholds that gave the top results in brackets.

### Distribution of the top clinical and gene features

The results of the two-sample KS-tests using the derivation set are shown in Table 3. Both the value of the statistic and the *p*-value is given for each feature. The higher value of the statistic corresponds to greater distributional differences. From this analysis, we can clearly see that the distributions of the top gene and clinical features have statistically significant differences between the complicated and uncomplicated course patients.

Using the same top 20 gene predictors, we repeated the two sample KS-tests on the validation set. However, in order to judge the sensitivity of the model towards different APACHE II scores, we tried different score cutoffs of 15, 20, 25, and 30. The list of genes that showed a significant distributional difference between the two classes and their corresponding KS scores are shown in Table 4. It can be seen from the analysis that an APACHE II cutoff of 25 gave the highest number (10) of significant genes showing that only a fraction of the top 20 biomarkers discovered using pediatric data is actually indicative of a complex course outcome in an adult cohort. We then performed repeated 5-fold cross-validation experiments on the validation data using these 10 genes and achieved the best results using the sampling technique SMOTE and XGBoost classifier pair (mean AUROC=0.72; mean Sensitivity=0.78; mean Specificity=0.65; mean MCC=0.4). These results were similar to the external validation results mentioned in the previous section. The Gaussian kernel density plots for the significant genes (*p*-value < 0.1) derived using both the derivation and validation data are shown in figure 7.

**Figure 7:**
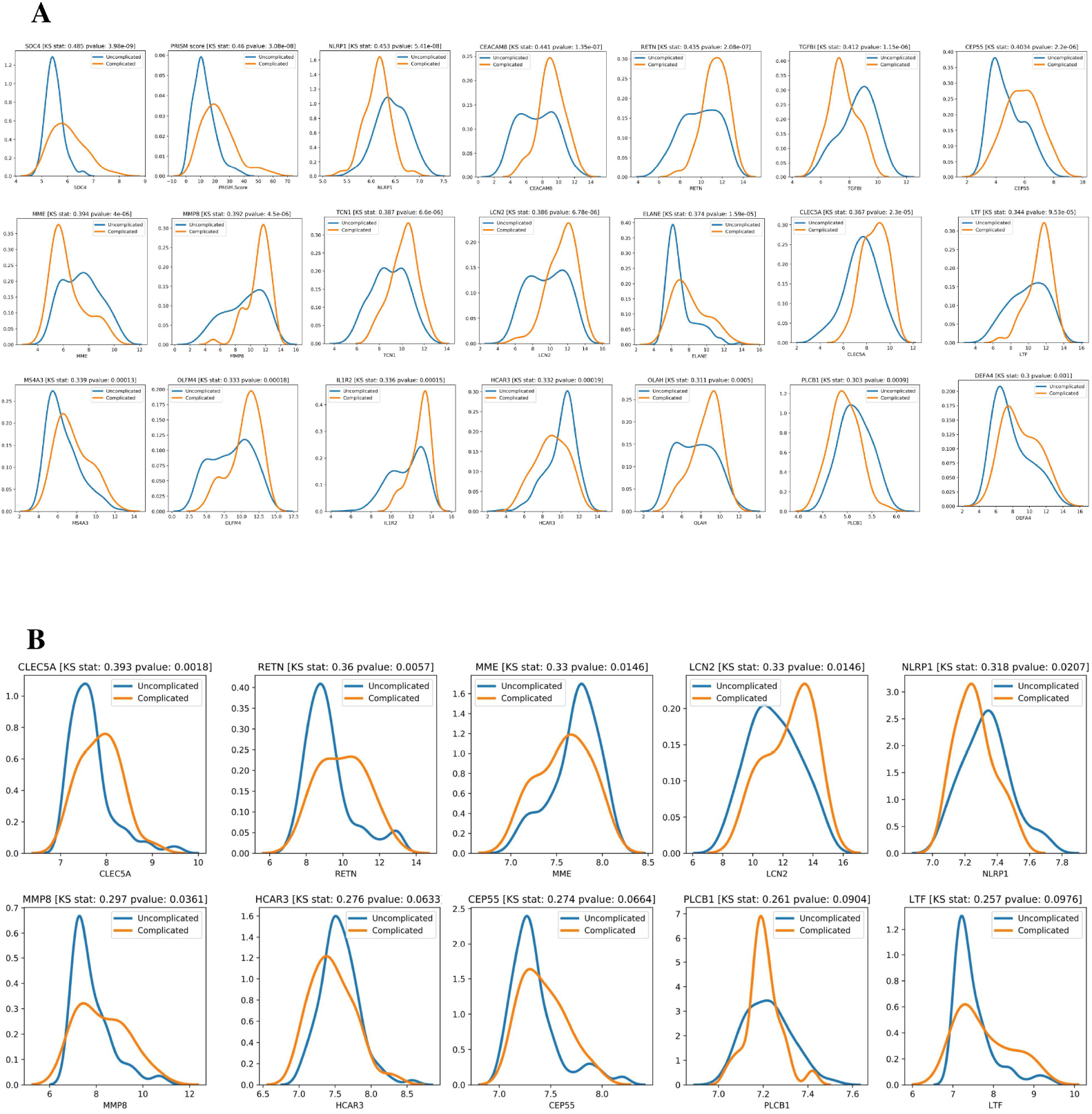
Class-wise gaussian kernel density plots for the top performing features along with the KS test scores. A Kolmogorov-Smirnov test is a non-parametric test used to compare the equality of probability distributions. There are two scores associated with a KS test: a KS statistic that is used to quantify the distance between two distributions and the p-value which tells us the significance of the result. (A) Distribution of the top 20 gene predictors and a severity score (PRISM) for the derivation dataset is shown in this plot (in decreasing order of KS statistic scores). (B) Out of the same top 20 gene predictors indicative of a complex course, only 10 gave significant KS statistic scores for the validation set which are shown in this figure.

**Table 4:**
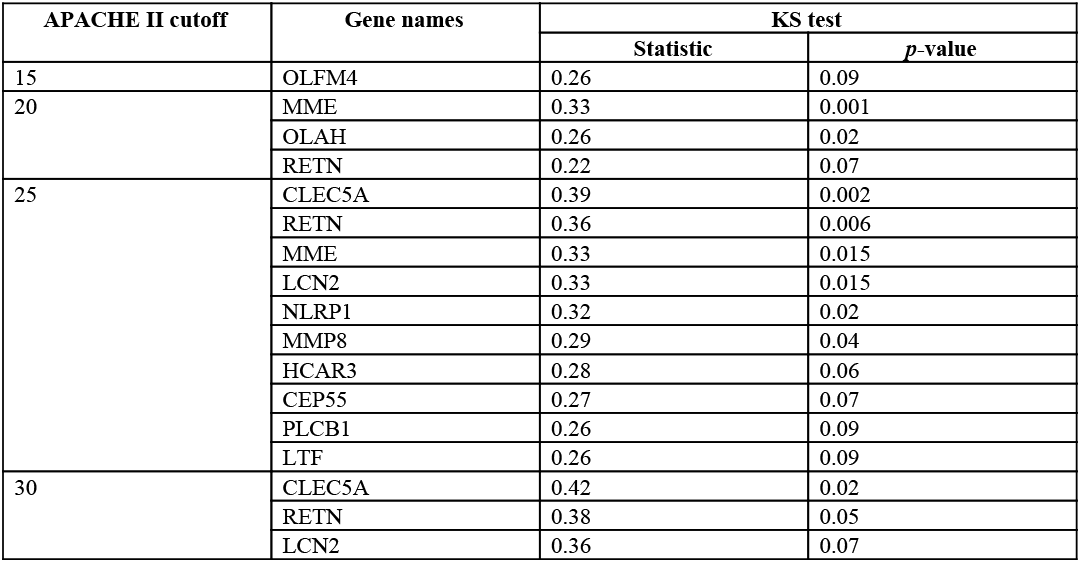
Number of significant genes (*P* < 0.1) obtained from the KS test analysis on the validation data using different APACHE II cutoffs

## Discussion

This study revealed differentially expressed genes in peripheral blood samples collected within 24 hours of PICU admission. Among the most significant genes from our machine learning analysis, we found genes that are responsible for innate immune response. Sepsis is caused by the dysregulation of the host response to an infection and in its severe form, causes life-threatening organ dysfunction. Cytokines play an important role during sepsis by regulating the immune response to the infection. A variety of pro-inflammatory and anti-inflammatory mediators contribute to the inflammatory response and an imbalance between the two directly results in dysregulation. Matrix metalloproteinase-8 (*MMP8*) and Resistin (*RETN*) have been associated with the activation of pro-inflammatory cytokines TNF-α (31, 32) which in turn stimulates systemic inflammation. *MMP8* is involved in the homeostasis of the extracellular space, largely expressed by mononuclear cells and macrophages; it has been shown to be involved in roles supporting innate immunity (33). *MMP8* knockout mice have been observed to have reduced phagocytosis and NET activity (34).

A number of the identified genes have been long implicated in Neutrophil Extracellular Traps (NETs), the activation of the NET results in NETosis, which can significantly complicate disease course. Among our genes, Olfactomedin 4 (*OLFM4*), Elastase Neutrophil Expressed (*ELANE*), NLR Family Pyrin Domain Containing 1 (*NLRP1*), lactotransferrin (*LTF*) were identified. Recent findings have suggested CEA Cell Adhesion Molecule 8 (*CEACAM8*), C-Type Lectin Domain Containing 5A (*CLEC5A*) (35) may be further implicated in NETosis. Polymorphonuclear neutrophils, typically pro-inflammatory, also perform immunoregulatory roles, by expressing *CEACAM8* and thus releasing soluble *CEACAM8* after activation. Extracellular chromatids have been shown to activate the secretion of *CEACAM8* through degranulation (36). A recent study finds *CLEC5A*, which has long been associated with dengue virus-induced lethal disease (37), to be an important factor associated with modulating innate immune response against bacterial infection in mice. That study suggests in mice with a knockout gene, that prognosis was severe and by day 5, the mice had significant liver necrosis and increased risk for death.

A combination of the genes identified in this analysis also has been involved in microbiome homeostasis. Specifically, Lipocalin-2 (*LCN2*), an innate immune protein has been associated with maintaining an intestinal barrier against oxidative stress, having immunosuppressive character, and protects against multi-organ dysfunction (38–40). This protein has been increasingly suggested to be a therapeutic candidate to protect against gut-origin sepsis (40).

The complexity of the disease may also contribute to the ambiguity in identifying the correct class of pathogens, specifically in gram-negative/gram-positive bacterial differentiation. Therefore, interest has emerged in differentiating these characteristics through gene expressions, and Interleukin 1 Receptor Type 2 (*IL1R2*) has been specifically implicated (41). Identifying such differentiation may also aid in determining the complexity and severity of sepsis by investigating the specific types of toxins released by either class of pathogens. Superantigens, for instance, produced by *S. aureus* and *S. pyrogenes* have been suggested to cause a massive cellular immune response, leading to fatal toxic shock (42), while other microbial toxins have been involved in significant sepsis-led immunosuppression (43). Hence, earlier identification of such differentiation can improve case management and therapeutic selection.

We identified Hydroxycarboxylic Acid Receptor 3 (*HCAR3*) and Membrane Metalloendopeptidase (*MME*) as two of the three downregulated genes similar to the findings of Kangelaris et al. (44) which studied changes in gene expression among ARDS (acute respiratory distress syndrome) patients affected with sepsis. The genes overexpressed in ARDS were often found to be associated with a more severe sepsis outcome in other studies (such as *MMP8, RETN, OLFM4*) (45, 46). Similar results were found among patients with Acute Kidney Injury (AKI) where the overexpression of genes such as *MMP8, IL1R2*, and *OLFM4* was associated with increased severity and organ failure (47).

Some of the genes identified in this work overlap with approaches to predict sepsis mortality (17). DEGs identified in our study namely, *CEACAM8, IL1R2, CEP55, TGFBI*, and *DEFA4* belonged to that list. Among the PERSEVERE genes (14) that were identified as DEGs in our analysis included *MMP8, ELA2, LCN2, LTF, andRETN* and those that overlapped with the PERSEVERE XP genes (15) included *CEP55, TGFBI, and MME*.

Among the top consistently chosen features (that were not DEGs) in our machine learning analysis, Syndecan-4 (*SDC4*) has been found to be a potential biomarker with an anti-inflammatory function in patients with acute pneumonia (48). Microbiologic results of the patients from our dataset, identified Pneumococcus as the second most frequently occurring pathogen hence this result is quite significant. We also identified Phospholipase C Beta 1 (*PLCB1*) as one of the top genes that were not a DEG. Phospholipase (*PLC*) proteins are mainly responsible for metabolizing inositol lipids and recent studies have shown that impairment of lipid metabolism is one of the major changes among patients with sepsis secondary to hospital-acquired pneumonia (49). Finally, the NLR Family Pyrin Domain Containing 1 (*NLRP1*) gene, which is one of the many genes responsible for the activation of the NLR-inflammasome cascades was found to be significantly altered among septic patients in a recent study (50).

The exclusion of patients meeting SIRS criteria resulted in a lower number of DEGs, and the machine learning prediction scores were also reduced, with an AUROC of 0.61. This implies that the inclusion of such patients within the non-complicated disease course resulted in a more robust characterization of complex patients, thereby improving the predictive performance. Moreover, five genes, namely *TGFBI, DEFA4, CEP55,MME, OLAH*, all except *CEP55* are implicated as early markers of neutrophil activity. *CEP55* overexpression has been implicated in T-cell lymphoma and genome instability (51, 52).

The gene expression profiles of the top 20 gene markers from the stability analysis had significant distributional differences between the complicated and uncomplicated course patients (Table 3, Figure 7a). When tested on a closely related independent adult validation set, we achieved an out-of-sample AUC of 0.72. The relatively lower AUC can be attributed to the fact that the derivation cohort is based on pediatric patients (mean age 3.81 yrs) while the validation cohort is based on adult patients (mean age 59.13 yrs). Some of the gene predictors that are indicative of a complex course outcome for pediatric patients, might not play the same role for adult patients. This was further justified when we found that out of the top 20 gene biomarkers, only 10 had a significant class-wise distributional difference using the gene expression profiles from the validation cohort using an APACHE II cutoff of 25 (Table 4, Figure 7b). Also, one of the consistently chosen top features from our analysis using the derivation data was the PRISM score which couldn’t be used for the validation analysis for obvious reasons. Secondly, even within the validation dataset, the complex disease course was interpreted based on severity of illness scores that were available at the time of ICU admission. Hence, a more similar dataset which derives complex disease course longitudinally during ICU stay may have improved the performance of the proposed biomarkers.

In this study, we identified a list of significant genes that were able to differentiate between sepsis patients with a complicated and uncomplicated course with reasonable accuracy. Blood-based biomarkers are minimally invasive and therefore can be used in conjunction with other clinical variables to detect complicated course outcomes among sepsis patients. There are a few limitations to our study. First, we present in this study a novel hybrid method of biomarker discovery that identifies stable and consistently chosen features among multiple iterations of the pipeline as candidate gene markers of complex disease course. While we try to balance this novel approach by demonstrating traditional methods of identifying DEGs, further investigations in other datasets may be required to establish its validity. Second, due to the lack of any similar public dataset that identifies complicated disease courses within sepsis patients, we identified a closely related dataset which presents severity of illness scores, using which we derived our interpretations of complex and noncomplex patients. Therefore, to ensure generalization and minimize selection bias, further validation needs to be performed on closely related prospective datasets. Thirdly, we derive our model in a pediatric cohort and validate it in an adult cohort. Finally, complicated and uncomplicated course outcomes were defined clinically and so there may be some limitations in that interpretation, which contribute bias.

## Conclusion

This paper presents a novel list of genes that predict a complicated disease course for critically ill patients in the pediatric intensive care unit. The list of 20 genes was derived from a rigorous feature selection and validation pipeline, wherein we measure feature stability over ten simulated iterations. Many of the derived genes were attributed to an innate immunity function and contributed to NETosis. The resulting derivation AUROC of 0.82 and validation AUROC of 0.723 suggests that these markers can reliably predict the outcome given only a single exam of peripheral blood. Future work can evaluate the performance of this gene list within a prospective trial.

## Acknowledgments

S.B. was supported by the Google Summer of Code 2019 program.

## Author Contributions

R.K. conceived and designed the study. S.B., R.K. and A.M. developed the method, performed the data analysis and interpretation. H.R.W. and N.P revised the article critically important for intellectual content. All authors wrote and proofread the manuscript.

## References

1. Severe Sepsis in the Emergency Department and Its Association With a Complicated Clinical Course - PubMed [Internet]. [cited 2020 May 30]. Available from: https://pubmed.ncbi.nlm.nih.gov/9864130/

2. Singer M, Deutschman CS, Seymour CW, Shankar-Hari M, Annane D, Bauer M, et al. The Third International Consensus Definitions for Sepsis and Septic Shock (Sepsis-3). JAMA. 2016 Feb 23;315(8):801.

3. Global, regional, and national sepsis incidence and mortality, 1990–2017: analysis for the Global Burden of Disease Study - The Lancet [Internet]. [cited 2020 May 30]. Available from: https://www.thelancet.com/journals/lancet/article/PIIS0140-6736(19)32989-7/fulltext

4. Seymour CW, Kennedy JN, Wang S, Chang C-CH, Elliott CF, Xu Z, et al. Derivation, Validation, and Potential Treatment Implications of Novel Clinical Phenotypes for Sepsis. JAMA. 2019 28;321(20):2003–17.

5. Leligdowicz A, Matthay MA. Heterogeneity in sepsis: new biological evidence with clinical applications. Critical Care. 2019 Mar 9;23(1):80.

6. Wong HR, Marshall JC. Leveraging Transcriptomics to Disentangle Sepsis Heterogeneity. Am J Respir Crit Care Med. 2017 Aug 1;196(3):258–60.

7. Long-term Host Immune Response Trajectories Among Hospitalized Patients With Sepsis | Critical Care Medicine | JAMA Network Open | JAMA Network [Internet]. [cited 2020 May 30]. Available from: https://jamanetwork.com/journals/jamanetworkopen/fullarticle/2747481

8. Islam MM, Nasrin T, Walther BA, Wu C-C, Yang H-C, Li Y-C. Prediction of sepsis patients using machine learning approach: A meta-analysis. Comput Methods Programs Biomed. 2019 Mar;170:1–9.

9. van Wyk F, Khojandi A, Mohammed A, Begoli E, Davis RL, Kamaleswaran R. A minimal set of physiomarkers in continuous high frequency data streams predict adult sepsis onset earlier. International Journal of Medical Informatics. 2019 Feb 1;122:55–62.

10. Kamaleswaran R, Akbilgic O, Hallman MA, West AN, Davis RL, Shah SH. Applying Artificial Intelligence to Identify Physiomarkers Predicting Severe Sepsis in the PICU. Pediatr Crit Care Med. 2018;19(10):e495–503.

11. The 2018 Surviving Sepsis Campaign’s Treatment Bundle: When Guidelines Outpace the Evidence Supporting Their Use - PubMed [Internet]. [cited 2020 May 30]. Available from: https://pubmed.ncbi.nlm.nih.gov/30193754/

12. Ibrahim ZM, Wu H, Hamoud A, Stappen L, Dobson RJB, Agarossi A. On classifying sepsis heterogeneity in the ICU: insight using machine learning. J Am Med Inform Assoc. 2020 Jan 17;27(3):437–43.

13. Engelen TSR van, Wiersinga WJ, Scicluna BP, Poll T van der. Biomarkers in Sepsis. Critical Care Clinics. 2018 Jan 1;34(1):139–52.

14. The Pediatric Sepsis Biomarker Risk Model - PubMed [Internet]. [cited 2020 May 30]. Available from: https://pubmed.ncbi.nlm.nih.gov/23025259/

15. Improved Risk Stratification in Pediatric Septic Shock Using Both Protein and mRNA Biomarkers. PERSEVERE-XP [Internet]. [cited 2020 May 30]. Available from: https://www.ncbi.nlm.nih.gov/pmc/articles/PMC5564676/

16. Mohammed A, Cui Y, Mas VR, Kamaleswaran R. Differential gene expression analysis reveals novel genes and pathways in pediatric septic shock patients. Sci Rep [Internet]. 2019 Aug 2 [cited 2020 May 30];9. Available from: https://www.ncbi.nlm.nih.gov/pmc/articles/PMC6677896/

17. Sweeney TE, Perumal TM, Henao R, Nichols M, Howrylak JA, Choi AM, et al. A community approach to mortality prediction in sepsis via gene expression analysis. Nature Communications. 2018 Feb 15;9(1):694.

18. Ready for Prime Time? Biomarkers in Sepsis - Emergency Medicine Clinics [Internet]. [cited 2020 May 30]. Available from: https://www.emed.theclinics.com/article/S0733-8627(16)30074-8/abstract

19. Sweeney TE, Shidham A, Wong HR, Khatri P. A comprehensive time-course-based multicohort analysis of sepsis and sterile inflammation reveals a robust diagnostic gene set. Sci Transl Med. 2015 May 13;7(287):287ra71.

20. Hyperchloremia Is Associated With Complicated Course and Mortality in Pediatric Patients With Septic Shock - PubMed [Internet]. [cited 2020 May 30]. Available from: https://pubmed.ncbi.nlm.nih.gov/29394222/

21. Parnell GP, Tang BM, Nalos M, Armstrong NJ, Huang SJ, Booth DR, et al. Identifying key regulatory genes in the whole blood of septic patients to monitor underlying immune dysfunctions. Shock. 2013 Sep;40(3):166–74.

22. affy—analysis of Affymetrix GeneChip data at the probe level | Bioinformatics | Oxford Academic [Internet]. [cited 2020 May 30]. Available from: https://academic.oup.com/bioinformatics/article/20/3/307/185980

23. gcrma: Background Adjustment Using Sequence Information version 2.58.0 from Bioconductor [Internet]. [cited 2020 May 30]. Available from: https://rdrr.io/bioc/gcrma/

24. Adjusting Batch Effects in Microarray Expression Data Using Empirical Bayes Methods - PubMed [Internet]. [cited 2020 May 30]. Available from: https://pubmed.ncbi.nlm.nih.gov/16632515/

25. limma powers differential expression analyses for RNA-sequencing and microarray studies | Nucleic Acids Research | Oxford Academic [Internet]. [cited 2020 May 30]. Available from: https://academic.oup.com/nar/article/43/7/e47/2414268

26. Yu G, Wang L-G, Han Y, He Q-Y. clusterProfiler: an R Package for Comparing Biological Themes Among Gene Clusters. OMICS. 2012 May;16(5):284–7.

27. Yu G, Hu E. enrichplot: Visualization of Functional Enrichment Result [Internet]. Bioconductor version: Release (3.11); 2020 [cited 2020 May 30]. Available from: https://bioconductor.org/packages/enrichplot/

28. Szklarczyk D, Gable AL, Lyon D, Junge A, Wyder S, Huerta-Cepas J, et al. STRING v11: protein–protein association networks with increased coverage, supporting functional discovery in genome-wide experimental datasets. Nucleic Acids Res. 2019 Jan 8;47(Database issue):D607–13.

29. PRISM III: An Updated Pediatric Risk of Mortality Score - PubMed [Internet]. [cited 2020 May 30]. Available from: https://pubmed.ncbi.nlm.nih.gov/8706448/

30. A stability index for feature selection | Proceedings of the 25th conference on Proceedings of the 25th IASTED International Multi-Conference: artificial intelligence and applications [Internet]. [cited 2020 May 30]. Available from: https://dl.acm.org/doi/10.5555/1295303.1295370

31. Silswal N, Singh AK, Aruna B, Mukhopadhyay S, Ghosh S, Ehtesham NZ. Human resistin stimulates the pro-inflammatory cytokines TNF-alpha and IL-12 in macrophages by NF-kappaB-dependent pathway. Biochem Biophys Res Commun. 2005 Sep 9;334(4):1092–101.

32. Matrix metalloproteinase-8 Plays a Pivotal Role in Neuroinflammation by Modulating TNF-α Activation - PubMed [Internet]. [cited 2020 May 31]. Available from: https://pubmed.ncbi.nlm.nih.gov/25049354/

33. Quintero PA, Knolle MD, Cala LF, Zhuang Y, Owen CA. Matrix metalloproteinase-8 inactivates macrophage inflammatory protein-1 alpha to reduce acute lung inflammation and injury in mice. J Immunol. 2010 Feb 1;184(3):1575–88.

34. Atkinson SJ, Varisco BM, Sandquist M, Daly MN, Klingbeil L, Kuethe JW, et al. Matrix metalloproteinase-8 augments bacterial clearance in a juvenile sepsis model. Molecular medicine (Cambridge, Mass). 2016 Aug 8;22:455–63.

35. CLEC5A Is a Critical Receptor in Innate Immunity Against Listeria Infection - PubMed [Internet]. [cited 2020 May 31]. Available from: https://pubmed.ncbi.nlm.nih.gov/28824166/

36. Frontiers | Extracellular Chromatin Triggers Release of Soluble CEACAM8 Upon Activation of Neutrophils | Immunology [Internet]. [cited 2020 May 31]. Available from: https://www.frontiersin.org/articles/10.3389/fimmu.2019.01346/full

37. CLEC5A Is Critical for Dengue-Virus-Induced Lethal Disease - PubMed [Internet]. [cited 2020 May 31]. Available from: https://pubmed.ncbi.nlm.nih.gov/18496526/

38. Moschen AR, Adolph TE, Gerner RR, Wieser V, Tilg H. Lipocalin-2: A Master Mediator of Intestinal and Metabolic Inflammation. Trends Endocrinol Metab. 2017;28(5):388–97.

39. Lipocalin 2 Protects From Inflammation and Tumorigenesis Associated With Gut Microbiota Alterations - PubMed [Internet]. [cited 2020 May 31]. Available from: https://pubmed.ncbi.nlm.nih.gov/27078067/

40. Lu F, Inoue K, Kato J, Minamishima S, Morisaki H. Functions and regulation of lipocalin-2 in gut-origin sepsis: a narrative review. Crit Care. 2019 Aug 2;23(1):269.

41. Lang Y, Jiang Y, Gao M, Wang W, Wang N, Wang K, et al. Interleukin-1 Receptor 2: A New Biomarker for Sepsis Diagnosis and Gram-Negative/Gram-Positive Bacterial Differentiation. Shock. 2017;47(1):119–24.

42. Superantigens: mechanism of T-cell stimulation and role in immune responses. - Abstract - Europe PMC [Internet]. [cited 2020 May 31]. Available from: https://europepmc.org/article/med/1832875

43. Gram-positive and gram-negative bacterial toxins in sepsis [Internet]. [cited 2020 May 31]. Available from: https://www.ncbi.nlm.nih.gov/pmc/articles/PMC3916377/

44. Kangelaris KN, Prakash A, Liu KD, Aouizerat B, Woodruff PG, Erle DJ, et al. Increased expression of neutrophil-related genes in patients with early sepsis-induced ARDS. Am J Physiol Lung Cell Mol Physiol. 2015 Jun 1;308(11):L1102–1113.

45. Chen Y, Shi J-X, Pan X-F, Feng J, Zhao H. DNA microarray-based screening of differentially expressed genes related to acute lung injury and functional analysis. Eur Rev Med Pharmacol Sci. 2013 Apr;17(8):1044–50.

46. Howrylak JA, Dolinay T, Lucht L, Wang Z, Christiani DC, Sethi JM, et al. Discovery of the gene signature for acute lung injury in patients with sepsis. Physiol Genomics. 2009 Apr 10;37(2):133–9.

47. Identification of Candidate Serum Biomarkers for Severe Septic Shock-Associated Kidney Injury via Microarray - PubMed [Internet]. [cited 2020 May 31]. Available from: https://pubmed.ncbi.nlm.nih.gov/22098946/

48. Serum Syndecan-4 as a Possible Biomarker in Patients With Acute Pneumonia - PubMed [Internet]. [cited 2020 May 31]. Available from: https://pubmed.ncbi.nlm.nih.gov/25895983/

49. Sharma NK, Ferreira BL, Tashima AK, Brunialti MKC, Torquato RJS, Bafi A, et al. Lipid metabolism impairment in patients with sepsis secondary to hospital acquired pneumonia, a proteomic analysis. Clin Proteomics. 2019;16:29.

50. Esquerdo KF, Sharma NK, Brunialti MKC, Baggio-Zappia GL, Assunção M, Azevedo LCP, et al. Inflammasome gene profile is modulated in septic patients, with a greater magnitude in non-survivors. Clinical & Experimental Immunology. 2017;189(2):232–40.

51. Expression and Clinical Significance of Centrosomal Protein 55 in T-cell Lymphoma - PubMed [Internet]. [cited 2020 May 31]. Available from: https://pubmed.ncbi.nlm.nih.gov/29516967/

52. Kalimutho M, Sinha D, Jeffery J, Nones K, Srihari S, Fernando WC, et al. CEP55 is a determinant of cell fate during perturbed mitosis in breast cancer. EMBO Mol Med [Internet]. 2018 Sep [cited 2020 May 31];10(9). Available from: https://www.ncbi.nlm.nih.gov/pmc/articles/PMC6127888/

